# Mechanism of nucleotide discrimination by the translesion synthesis polymerase Rev1

**DOI:** 10.1101/2021.12.01.470840

**Authors:** Tyler M. Weaver, Timothy H. Click, Thu H. Khoang, M. Todd Washington, Pratul K. Agarwal, Bret D. Freudenthal

**Affiliations:** Department of Biochemistry and Molecular Biology, Department of Cancer Biology, University of Kansas Medical Center, Kansas City, Kansas, 66160; Department of Physiological Sciences and High-Performance Computing Center, Oklahoma State University, Stillwater, Oklahoma 74048; Department of Biochemistry, University of Iowa, Iowa City, IA 52242

**Keywords:** DNA polymerase, DNA repair, Translesion synthesis

## Abstract

Rev1 is a translesion DNA synthesis (TLS) polymerase involved in the bypass of adducted-guanine bases and abasic sites during DNA replication. During damage bypass, Rev1 utilizes a protein-template mechanism of DNA synthesis, where the templating DNA base is evicted from the Rev1 active site and replaced by an arginine side chain that preferentially binds incoming dCTP. Here, we utilize X-ray crystallography and molecular dynamics simulations to obtain structural insight into the dCTP specificity of Rev1. We show the Rev1 R324 protein-template forms sub-optimal hydrogen bonds with incoming dTTP, dGTP, and dATP that prevents Rev1 from adopting a catalytically competent conformation. Additionally, we show the Rev1 R324 protein-template forms optimal hydrogen bonds with incoming rCTP. However, the incoming rCTP adopts an altered sugar pucker, which prevents the formation of a catalytically competent Rev1 active site. This work provides novel insight into the mechanisms for nucleotide discrimination by the TLS polymerase Rev1.

## Introduction

DNA polymerases are tasked with the faithful replication of genomic DNA during each cycle of cell division^1–3^. The faithful replication of the genome requires DNA polymerases bind and add the correct incoming deoxynucleotide triphosphate (dNTP) to the end of a growing DNA primer strand. Importantly, the correct nucleotide must be chosen amongst a pool of excess incorrect nucleotides that contain either the wrong DNA nucleobase (i.e. base) or the wrong sugar moiety (e.g. ribonucleotide triphosphates, rNTP). DNA polymerases use an induced-fit mechanism for selecting and incorporating the correct dNTP^4–8^. In the induced-fit mechanism, the DNA polymerase undergoes a conformational change during dNTP binding, which allows for sampling of proper Watson-crick base pairing with the templating DNA base. When the correct nucleotide is bound the 3′-OH of the primer terminus, Pα of the incoming nucleotide, and catalytic metal are in the proper orientation for catalysis within the active site^9^. If the incorrect nucleotide is bound, the sub-optimal base pairing between the incoming nucleotide and templating DNA base leads to conformational changes in the active site that are not conducive to catalysis^9, 10^. In addition to DNA base fidelity, DNA polymerases also possess mechanisms to maintain sugar fidelity. This includes an active site bulky aromatic steric gate residue that clashes with the 2′-OH of the incoming ribonucleotide triphosphate^11, 12^. In some cases, DNA polymerases also use a polar filter residue that locks the C2′ of the incoming ribonucleotide near the steric gate residue^13^. Ultimately, the ability to bind the correct incoming nucleotide enables DNA polymerases to faithfully replicate and repair genomic DNA.

Rev1 is a specialized Y-family DNA polymerase involved in translesion DNA synthesis (TLS), replication of G4-quadruplex DNA^14–16^, replication-induced gap filling^17^, and somatic hypermutation^18^. Rev1 is best known for its functions during TLS, where it acts as both a scaffolding component and a TLS DNA polymerase^19–23^. The DNA synthesis activity of Rev1 is important for the bypass of adducted-guanine bases and abasic sites, which otherwise stall replicative DNA polymerases and potentially collapse replication forks. The initial characterization of the Rev1 DNA synthesis activity observed a significant preference for cytosine insertion, irrespective of the templating DNA base identity^24–28^. Subsequent structural studies revealed the preference for cytosine insertion is due to the use of an unusual protein-template mechanism^29–34^. In the protein-template mechanism, the templating DNA base is evicted from the Rev1 active site and is replaced by an arginine side chain residue (Fig. 1). This arginine side chain forms two hydrogen bonds with an incoming dCTP, which is thought to enable the preferentially insertion of dCTP. Notably, Rev1 is the only DNA polymerase known to use an amino-acid side chain to encode for an incoming nucleotide, instead of a templating DNA base^2^. Additionally, Rev1 does not undergo large conformational changes upon incoming nucleotide binding. This makes Rev1 an interesting system to study mechanisms of nucleotide discrimination by a non-classical DNA polymerase.

**Figure 1.**
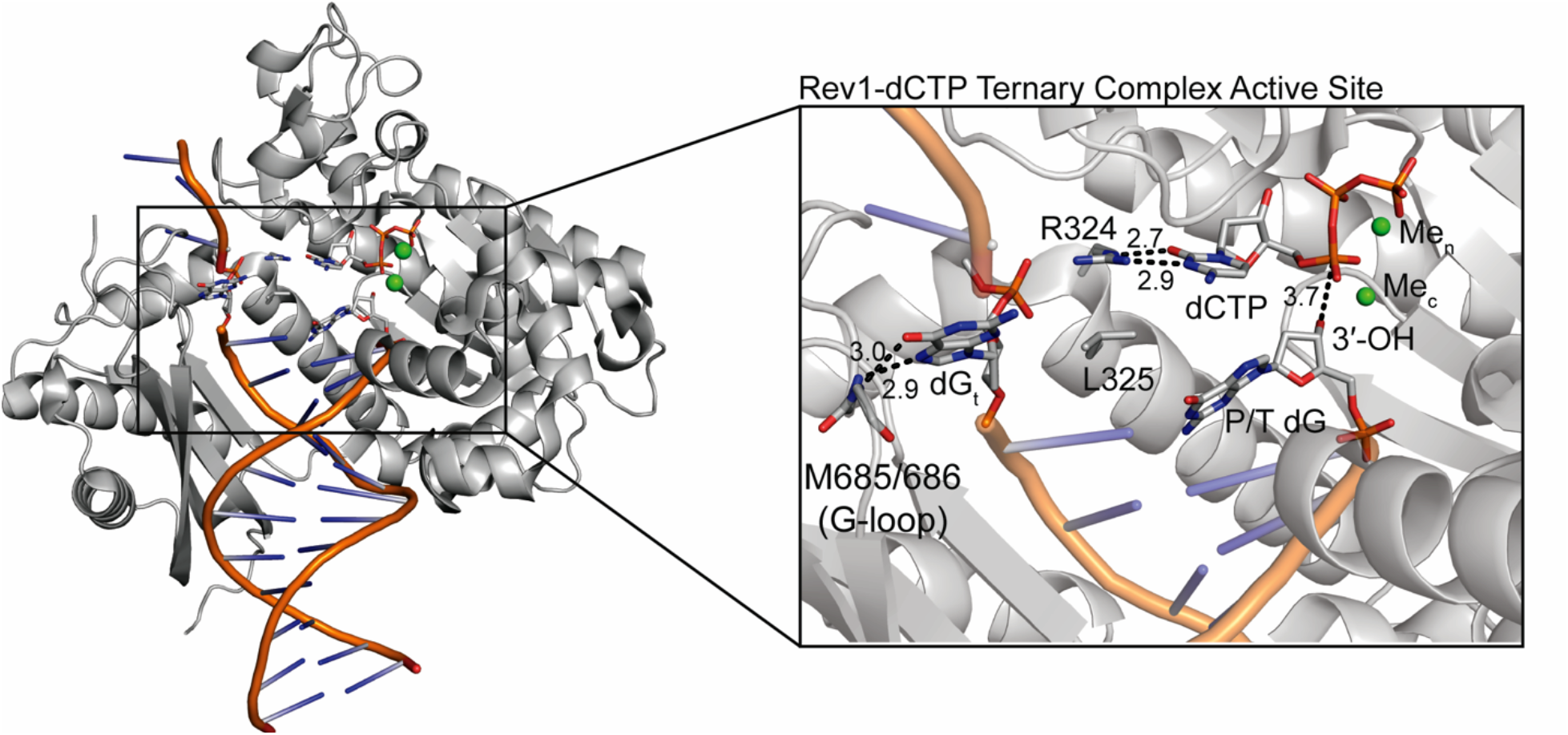
Rev1/DNA/dCTP ternary complex structure. An overall structure (left) and active site closeup (right) of the Rev1/DNA/dCTP ternary complex (PDB:6X6Z). Key protein and DNA residues are shown as grey sticks and labeled. The nucleotide binding (Me_n_) and catalytic (Me_c_) metals are shown as green spheres.

Several groups have studied the base and sugar discrimination of Rev1 using enzyme kinetics^24–28, 35, 36^. With respect to base discrimination, human Rev1 preferentially incorporates a dCTP opposite a template G^36, 37^. dGTP and dTTP are incorporated opposite G with 260- and 140-fold reduced relative catalytic efficiency (k_pol_/K_d_), respectively, relative to dCTP incorporation^37^. Rev1 incorporates dATP opposite a template G with a 14,000-fold reduced relative catalytic efficiency (k_pol_/K_d_) relative to dCTP incorporation^37^. With respect to sugar discrimination, Rev1 preferentially incorporations rCTP opposite a templating G with a 280-fold reduced relative catalytic efficiency (k_pol_/K_d_) compared to dCTP^37^. The structural basis for the dCTP specificity of Rev1 has been unclear, largely owing to the absence of structural information about the enzyme-substrate complexes with sub-optimal incoming NTPs. Here, we utilize X-ray crystallography and molecular dynamics (MD) simulations to obtain insight into the structure and dynamics of Rev1 interacting with different incoming NTPs, which explains the structural basis for Rev1 dCTP specificity.

## Results

### Structural basis for Rev1 DNA base specificity

To obtain structural insight for the interaction of Rev1 and an incoming dTTP, we determined a ternary complex (Rev1/DNA/dNTP) structure of Rev1 with an incoming dTTP. To accomplish this, Rev1 was crystallized in complex with a primer-template DNA generating Rev1/DNA binary complex crystals. The Rev1/DNA binary complex crystals were then transferred to a cryoprotectant containing dTTP and CaCl_2_ to generate Rev1-dTTP ternary complex crystals. Importantly, Ca^2+^ permits incoming nucleotide binding but does not facilitate catalysis *in crystallo*^34^. The resulting Rev1/DNA/dTTP ternary complex crystal diffracted to 1.70 Å and was in space group P2_1_2_1_2_1_ (Supplementary Table 2).

The overall structure of the Rev1-dTTP ternary complex is similar to the Rev1-dCTP ternary complex structure (RMSD = 0.228 Å), including the evicted templating guanine base, the R324 protein template, and Rev1 active site side chains (Supplementary Fig. 1). However, major differences were seen in the conformation of the incoming dTTP and the primer terminal dG DNA base (Supplementary Fig. 1 and Figure 2). Figure 2a shows the overall Rev1-dTTP ternary complex structure and a close-up of the Rev1 active site. The Rev1-dTTP ternary complex contains the incoming dTTP and both nucleotide (Ca_n_) and catalytic (Ca_c_) calcium ions (Fig. 2a). The incoming dTTP is coordinated by the R324 through a single non-planar hydrogen bond between the O2 of the Watson-Crick edge and the Nε of R324 (Fig. 2b). Notably, the position of the dTTP ribose sugar and triphosphate group are in a similar conformation to that seen in the Rev1-dCTP ternary complex structure (Fig. 2c). Structural superimposition of the Rev1-dTTP and Rev-dCTP ternary complex structures revealed several changes to the Rev1 active site with implications for dTTP insertion (Fig. 2c). The non-planar R324-dTTP interaction leads to a 2.6 Å movement in the dTTP base towards the primer terminal dG. To accommodate the dTTP, the primer terminal dG moves 1.4 Å downward from the conformation seen in the Rev1-dCTP ternary complex structure. This brings the 3′-OH of the primer terminal dG 4.4 Å away from Pα of the incoming dTTP, which is ∼1 Å increase compared to the Rev1-dCTP ternary complex structure (Fig. 2c, Fig. 1a). This increase shifts the 3’-OH out of a catalytically competent conformation to support nucleophilic attack on the Pα of the incoming dTTP. Ultimately, the loss of a single hydrogen bond between R324 and the dTTP and the altered conformation of the primer terminal dG 3’-OH explains the observed reduction in nucleotide binding and catalysis that was previously described for Rev1 insertion of dTTP (Supplementary Table 1)^36, 37^.

**Figure 2.**
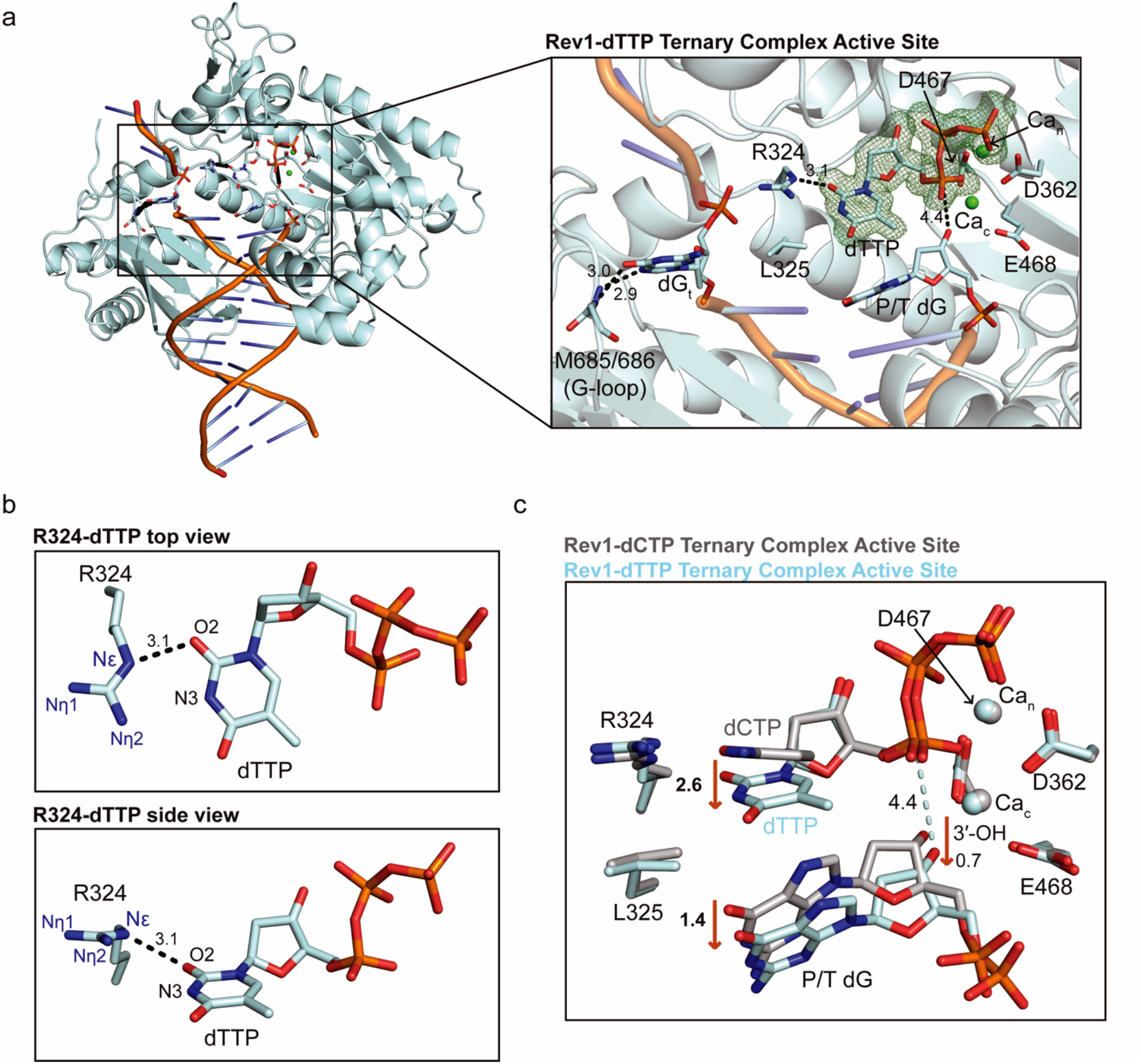
Rev1/DNA/dTTP ternary complex structure. **(a)** An overall structure (left) and active site closeup (right) of the Rev1/DNA/dTTP ternary complex. Key protein and DNA residues are shown as blue sticks. A Polder OMIT map contoured at σ=3.0 for the incoming dTTP is shown as a green mesh. **(b)** Two focused views of the hydrogen bonds formed between Rev1 R324 and the incoming dTTP. Hydrogen bonds are shown as black dashed lines and hydrogen bonding distances are labeled. **(c)** An overlay of the Rev1/DNA/dTTP ternary complex (light blue sticks) and Rev1/DNA/dCTP ternary complex (grey sticks). Red arrows and the respective distances (Å) highlight key active site differences between the two structures.

To obtain structural insight into Rev1 interacting with an incoming dGTP, we determined a 2.0 Å structure of the Rev1/DNA/dGTP ternary complex (Supplementary Table 2). The overall Rev1-dGTP ternary complex structure is very similar to the Rev1-dCTP ternary complex (RMSD = 0.306 Å, Supplementary Fig. 2). However, major alterations to the Rev1 active site were observed including the conformation of the dGTP and subtle conformational changes in the primer terminal DNA base were observed (Supplementary Fig. 2 and Figure 3). Figure 3a shows the overall Rev1-dGTP ternary complex structure and a close-up of the Rev1 active site, which contains the incoming dGTP and nucleotide Ca_n_ ion. Interestingly, the catalytic Ca_c_ was not observed and the Ca_c_ binding site was taken up by a water molecule. In the Rev1-dGTP ternary complex structure, the dGTP base adopts a Hoogsteen conformation, instead of the traditional Watson-Crick conformation. In this conformation, the Hoogsteen edge of the dGTP faces the R324 protein-template. The Nε and Nη of R324 form two non-planar hydrogen bonds with the N7 and O6 of the dGTP Hoogsteen edge (Fig. 3b). Interestingly, the dGTP ribose adopts a non-canonical C1’-exo sugar pucker, which is different than the C3’-endo sugar pucker seen with incoming dCTP (Fig. 3a and 3c). Despite the differences in DNA base and sugar conformations, the triphosphate group of the dGTP is anchored in the Rev1 active site in similar conformation as incoming dCTP (Fig. 3c and 3d). Structural superimposition of the Rev1-dGTP and Rev1-dCTP ternary complex structures revealed additional changes to the Rev1 active site with implications for catalysis (Fig. 3c and 3d). The non-planar interactions between the dGTP and R324 lead to a 1.9 Å movement of the dGTP base towards the primer terminal dG. This conformational change leads to a subtle 0.8 Å movement in the 3’-OH of the primer terminal dG distal to the Pα of the incoming dGTP (Fig. 3D). In this conformation, the 3’-OH is 4.4 Å from the Pα of the incoming dGTP, which is slightly longer than the 3.7 Å seen in the Rev1-dCTP ternary complex (Fig. 3a). Ultimately, the Hoogsteen conformation of the dGTP explains the reduction in nucleotide binding, while the subtle movement in the 3’OH distal to the Pα can explain the minor decrease in catalysis that was previously described for Rev1 insertion of dGTP (Supplementary Table 1)^36, 37^.

**Figure 3.**
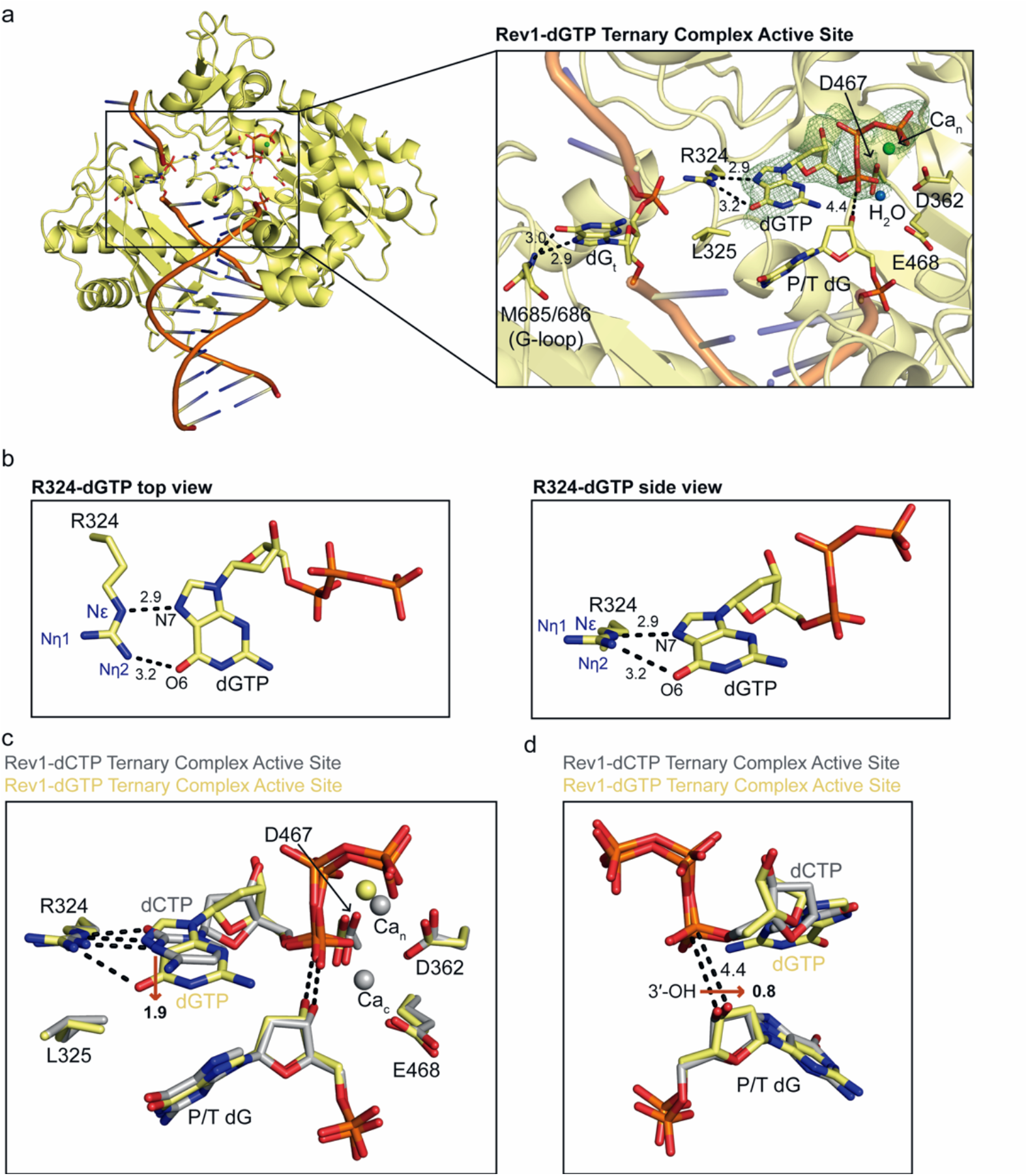
Rev1/DNA/dGTP ternary complex structure. **(a)** An overall structure (left) and active site closeup (right) of the Rev1/DNA/dGTP ternary complex. Key protein and DNA residues are shown as yellow sticks. A Polder omit map contoured at σ=3.0 for the incoming dGTP is shown as a green mesh. **(b)** Two focused views of the hydrogen bonds formed between Rev1 R324 and the incoming dGTP. Hydrogen bonds are shown as black dashed lines and hydrogen bonding distances (Å) are labeled. **(c)** An overlay of the Rev1/DNA/dGTP ternary complex (yellow sticks) and Rev1/DNA/dCTP ternary complex (grey sticks) showing differences in the dNTP bases. Red arrows and distances (Å) highlight key differences between the two structures. **(d)** An overlay of the Rev1/DNA/dGTP ternary complex (yellow sticks) and Rev1/DNA/dCTP ternary complex (grey sticks) showing differences in the position of the primer terminal dG 3’-OH. Red arrows and the respective distances (Å) show the movement of the primer terminal dG 3’-OH between the two structures.

To understand how Rev1 interacts with incoming dATP, we determined a 1.8 Å structure of the Rev1/DNA/dATP ternary complex (Supplementary Table 2). The overall structure of the Rev1-dATP ternary complex is very similar to the Rev1-dCTP ternary complex, including the evicted templating guanine base, the R324 protein template, and the important Rev1 active site side chains (RMSD=0.331, Supplementary Fig. 3). However, major alterations to the conformation of the dATP and the primer terminal DNA base were observed(Supplementary Fig. 3 and Figure 4). Figure 4a shows a close-up of the Rev1-dATP ternary complex active site, which contains the incoming dATP and both Ca_n_ and Ca_c_ ions. Interestingly, we observed two dATP conformations in the Rev1 active site. In both conformations, the dATP base adopts the Hoogsteen conformation, with the Hoogsteen edge facing the R324 protein-template. In conformation 1, the dATP forms a single non-planar hydrogen bond between the N7 of the Hoogsteen edge and the Nε of R324 (Fig. 4b). In conformation 2, the dATP forms a single non-planar hydrogen bond between the N7 of the dATP Hoogsteen edge and the Nε of R324, but the incoming dATP has undergone a 1.7 Å downward movement in relation to R324 that was not seen in dATP conformation 1 (Fig. 4c). Additional differences in the conformation of the dATP ribose sugar were observed. In the dATP conformation 1, the ribose sugar adopts a C4’-exo sugar pucker. (Fig. 4d). In the dATP conformation 2, the ribose sugar adopts a C2’-endo sugar pucker (Fig. 4e). Notably, the sugar puckers for both dATP conformations are different than the C3’-endo sugar pucker observed for incoming dCTP. Structural superimposition of the Rev1-dATP and Rev1-dCTP ternary complex structures identified additional changes within the active site with implications for catalysis (Fig. 4d and e). The non-planar interactions of both dATP conformation 1 and 2 leads to a 1.1 Å movement of the primer terminal dG 5′-phosphate, which results in a 1.0 Å movement of the 3’-OH distal to the Pα of the incoming dATP. This brings the 3’-OH to 4.0 Å and 4.4 Å from Pα of the dATP in conformation 1 and 2, respectively (Fig. 4D and 4E). In both conformations, the 3’-OH is not in the proper conformation for in-line nucleophilic attack on Pα of the incoming dATP. Together, this indicates that the Hoogsteen conformation of the dATP leads to the reduction in nucleotide binding, while the movement in the 3’-OH distal to the Pα explains the decrease in catalysis that was previously described for Rev1 insertion of dATP (Supplementary Table 1)^36, 37^.

**Figure 4.**
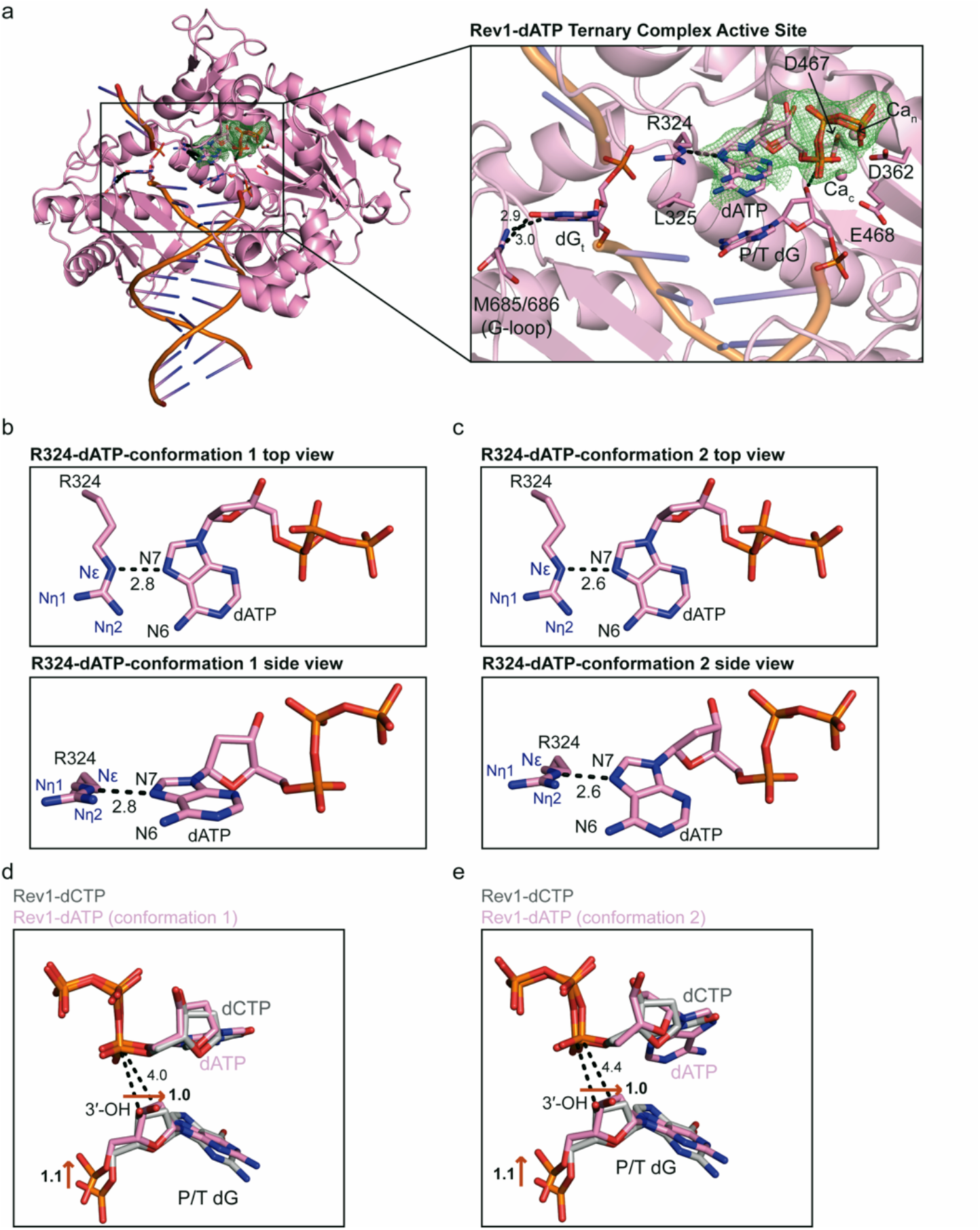
Rev1/DNA/dATP ternary complex structure. **(a)** An overall structure (left) and active site closeup (right) of the Rev1/DNA/dATP ternary complex. Key protein and DNA residues are shown as pink sticks. A Polder OMIT map contoured at σ=3.0 for the incoming dATP is shown as a green mesh. **(b)** Two focused views of the hydrogen bonds formed between Rev1 R324 and the incoming dATP (conformation 1). Hydrogen bonds are shown as black dashed lines and hydrogen bonding distances (Å) are labeled. **(c)** Two focused views of the hydrogen bonds formed between Rev1 R324 and the incoming dATP (conformation 2). Hydrogen bonds are shown as black dashed lines and hydrogen bonding distances (Å) are labeled. **(d)** An overlay of the Rev1/DNA/dATP ternary complex (conformation 1, pink sticks) and Rev1/DNA/dCTP ternary complex (grey sticks) showing differences in the position of the primer terminal dG 3’-OH. Red arrows and the respective distances (Å) show the movement of the primer terminal dG 3’-OH between the two structures. **(e)** An overlay of the Rev1/DNA/dATP ternary complex (conformation 2, pink sticks) and Rev1/DNA/dCTP ternary complex (grey sticks) showing differences in the position of the primer terminal dG 3’-OH. Red arrows and the respective distances (Å) show the movement of the primer terminal dG 3’-OH between the two structures.

### Dynamics of the Rev1-dNTP ternary complex active sites from MD simulations

Our structural analysis reveals that Rev1 can bind an incoming dTTP, dGTP, and dATP. However, these structures represent static snapshots of the incoming nucleotide in the Rev1 active site. To further understand the stability and dynamics of the R324 interaction with different incoming dNTPs, MD simulations were performed on the Rev1-dTTP, Rev1-dATP, and Rev1-dGTP ternary complex structures. We previously performed 1 μs MD simulations on the Rev1-dCTP ternary complex, which provided insight into the stability and dynamics of the Rev1 active site. In the Rev1-dCTP ternary complex simulation, we were able to monitor stable hydrogen bonds between the O2-Nε (100% of snapshots) and the N3-Nη (99.9% of snapshots) throughout the trajectory of the MD simulation (Supplementary Fig. 4). Therefore, these MD simulations can provide critical insight into the interaction of R324 with each different incoming dNTP base.

For this study, we initially performed a 1 μs MD simulation on the Rev1-dTTP ternary complex and monitored distances between the O2 and N3 of dTTP and the Nε and Nη of the R324 guanidinium group. During the MD simulation, we observed hydrogen bonding between the O2 of the dTTP and the Nε of R324 during 19.7% of the structural snapshots. In contrast, the N3 of the dTTP was within hydrogen bonding distance of the R324 Nη of R324 during only 0.4% of the MD snapshots, which is likely due to repulsion between the protonated dTTP N3 and protonated R324 Nη (Fig. 5a). This indicates the interaction of the dTTP base with R324 is largely driven by the single O2-Nε hydrogen bond, which is consistent with the X-ray crystal structure of the Rev-dTTP ternary complex (Fig. 2a). Interestingly, this data also suggests that the R324-dTTP interaction is dynamic on the ns-μs timescale.

**Figure 5.**
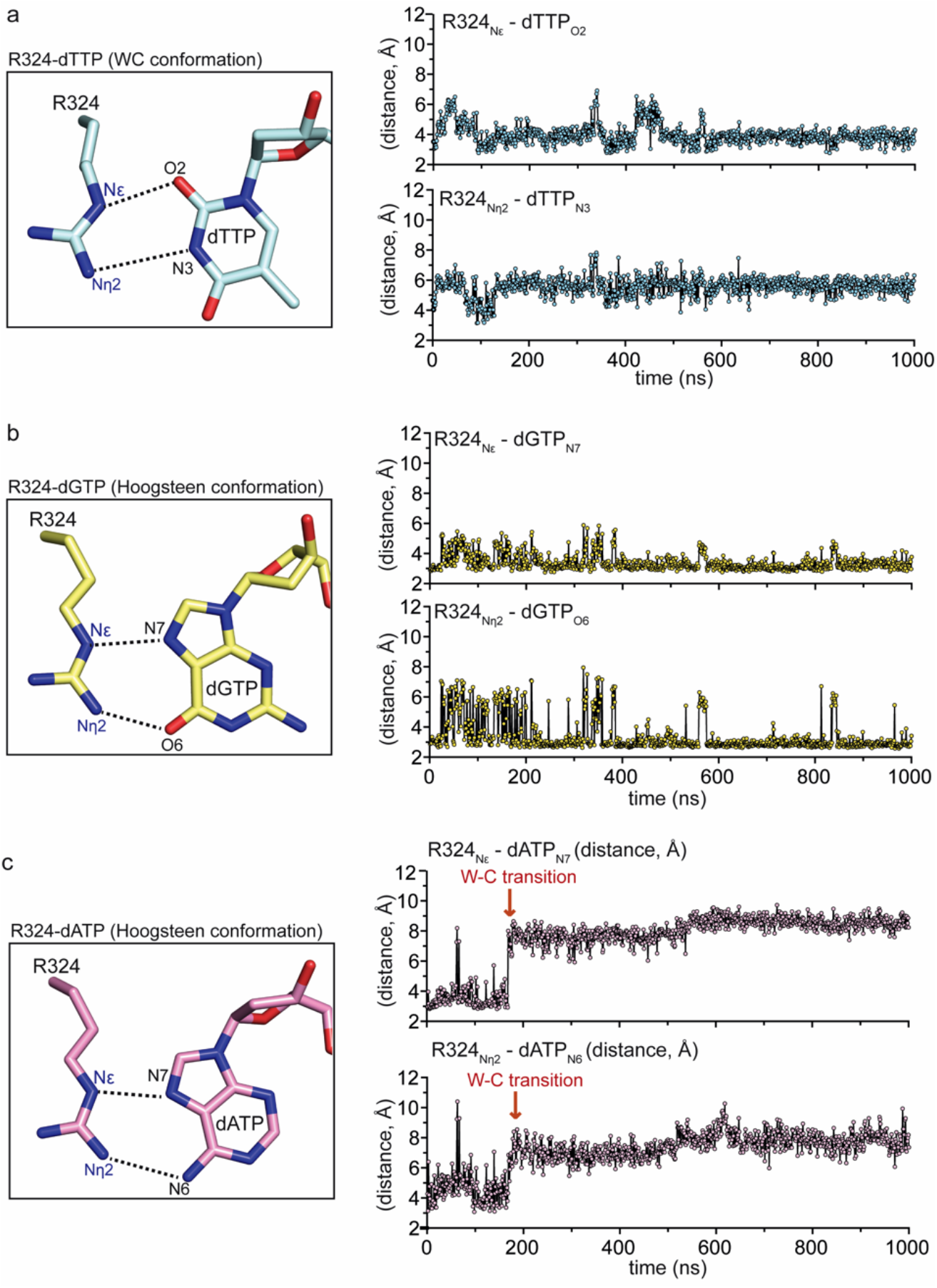
Molecular dynamics simulations of Rev1/DNA/dNTP ternary complex structures. **(a)** Focused view of the Rev1 R324 and incoming dTTP (Watson-crick conformation) showing the distances monitored throughout the MD simulation (left). Distance profiles for R324_NE_-dTTP_O2_ and the R324_Nn2_-dTTP_N3_ in the Rev1/DNA/dTTP ternary complex simulation (right). Each datapoint represents the distance (Å) between the indicated atoms at a single snapshot (1ns) from the MD simulation. **(b)** Focused view of the Rev1 R324 and incoming dGTP (Hoogsteen conformation) showing the distances monitored throughout the MD simulation (left). Distance profiles for R324_NE_-dGTP_N7_ and the R324_Nn2_-dGTP_O6_ in the Rev1/DNA/dGTP ternary complex simulation (right). Each datapoint represents the distance (Å) between the indicated atoms at a single snapshot (1ns) from the MD simulation. **(c)** Focused view of the Rev1 R324 and incoming dATP (Hoogsteen conformation) showing the distances monitored throughout the MD simulation (left). Distance profiles for R324_NE_-dATP_N7_ and the R324_Nn2_-dATP_N5_ in the Rev1/DNA/dATP ternary complex simulation (right). Each datapoint represents the distance (Å) between the indicated atoms at a single snapshot (1ns) from the MD simulation. The dATP transition from Hoogsteen to Watson-Crick conformation is denoted with a red arrow.

We also performed a 1 μs MD simulation on the Rev1-dGTP ternary complex and monitored the distance between the N7 and O6 of the dGTP Hoogsteen edge and the Nε and Nη of the R324 guanidinium group (Fig. 3b). During the MD simulation, hydrogen bonding was observed between the N7 of the dGTP Hoogsteen edge and the Nε during 73.4% of the MD trajectory (Fig. 5b). Similarly, hydrogen bonding was also observed between O6 of the dGTP Hoogsteen edge and Nη of R324 during 79.1% of the MD trajectory (Fig. 5b). This indicates that the interaction between the dGTP Hoogsteen edge and the R324 protein template is driven by the N7-Nε and O6-Nη hydrogen bonds, and that these hydrogen bonds are stable in the Rev1 active site (Fig. 3b).

We subsequently performed a 1 μs MD simulation on the Rev1-dATP ternary complex and monitored the distance between the N7 and N6 of the dATP Hoogsteen edge and the Nε and Nη of the Rev1 R324 protein-template. During the simulation, hydrogen bonding between the N7 of the dATP and Nε of Rev1 was observed during 9.8% of the MD trajectory. Similarly, the N6 of the dATP is within hydrogen bonding distance of the Rev1 Nη during only 2.2% of the MD trajectory. The minimal snapshots of dATP N6 within hydrogen bonding distance of Rev1 Nη is likely due to repulsion between the two protonated atoms. This indicates that the Hoogsteen edge of the dATP does not form a stable interaction with the R324 protein template. Interestingly, after ∼200 ns we observed a significant conformational change in the dATP base that corresponds to a switch from the Hoogsteen conformation to the Watson-Crick conformation (Fig. 5c). The dATP Watson-Crick face has a single hydrogen bond acceptor (dATP N1) that could potentially interact with the Nε or Nη of R324. However, an interaction between the dATP N1 and the Nε or Nη of R324 was not observed during the 800 ns of the simulation that the dATP adopts the Watson-Crick conformation (Supplementary Fig. 5a). Finally, we carried out a 1 μs MD simulation with the dATP in a Watson-Crick starting conformation to determine if an interaction with R324 could be observed. Notably, a stable interaction between dATP and R324 was also not observed in this simulation (Supplementary Fig. 5b). These MD simulations are consistent with dATP being highly dynamic in the Rev1 active site, which is due to a lack of stable hydrogen bonds between R324 and the Hoogsteen- or Watson-Crick edge of the dATP base.

### Structural basis for Rev1 sugar selectivity

In addition to DNA base discrimination, DNA polymerases must also maintain sugar selectivity to preferentially incorporate dNTPs rather than rNTPs. This is generally accomplished through a steric gate and/or polar filter residue that prevents ribonucleotide binding and incorporation. Despite the presence of both a steric gate and polar filter residue, Rev1 displays poor sugar selectivity inserting dCTP with a relative catalytic efficiency that is 280-fold higher than rCTP (Supplementary Table 1)^37^. Notably, this sugar selectivity is orders of magnitude worse than most DNA polymerases^11, 12^. To obtain mechanistic insight into the modest sugar selectivity by Rev1, we determined a 1.75 Å structure of the Rev1/DNA/rCTP ternary complex (Supplementary Table 2).

The overall Rev1-rCTP ternary complex structure is very similar to the Rev1-dCTP ternary complex structure (RMSD = 0.258 Å), including the evicted templating guanine base, the R324 protein template, and important Rev1 active site side chains (Supplementary Fig. 6a). However, we observed significant differences in the conformation of the incoming rCTP sugar and the primer terminal DNA base. Figure 6a shows a close-up of the active site for the Rev1-rCTP ternary complex, which contains the incoming rCTP and both Ca_n_ and Ca_c_ ions. The rCTP base forms two planar hydrogen bonds with the R324 protein template utilizing the N2 and O6 of the Watson-Crick face and the Nε and Nη of the R324 guanidinium group (Fig. 6b). This interaction of the rCTP with R324 is very similar to the interaction of the incoming dCTP and R324 (Fig. 1a, Supplementary Fig. 6a). Notably, a significant conformational change was observed in the ribose sugar of the rCTP, which adopts a C2’-endo sugar pucker compared to the C3’-endo sugar pucker observed for incoming dCTP (Fig. 6c, Supplementary Fig. 6b). In this conformation, the closest distance between the 2’-OH and F367 is 3.2 Å, indicating the 2’-OH does not significantly clash with the Rev1 F367 steric gate (Fig. 6c). In contrast, if the rCTP was in a C3’-endo sugar pucker the 2’-OH would have significant clashes with the F367 steric gate (Fig. 6c). Rev1 also possesses a polar filter residue (S402), which is thought to interact with the 3’-OH of the incoming nucleotide and position the C2’ in close proximity to the steric gate (Supplementary Fig. 6c)^13^. This interaction would provide an additional layer of discrimination for dCTP over rCTP. Interestingly, the C2’-endo sugar pucker leads to a conformation of the rCTP 3’-OH that does not interact with the S402 polar filter residue (Supplementary Fig. 6c). Taken together, the C2’-endo sugar pucker of the rCTP prevents significant clashing with Rev1, thus enabling the active site to readily accommodate insertion of ribonucleotides.

**Figure 6.**
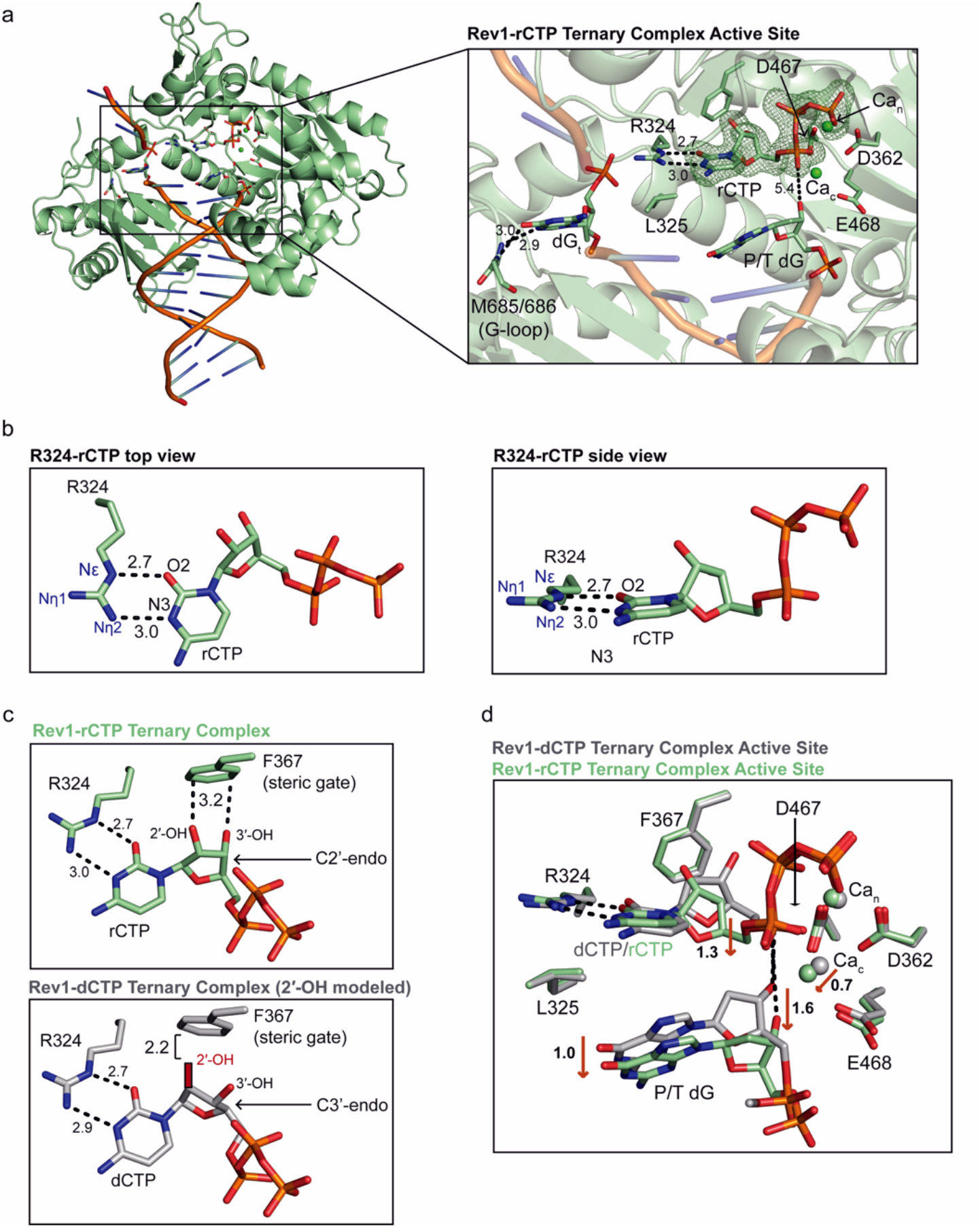
Rev1/DNA/rCTP ternary complex structure. **(a)** An overall structure (left) and active site closeup (right) of the Rev1/DNA/rCTP ternary complex. Key protein and DNA residues are shown as green sticks. A Polder OMIT map contoured at σ=3.0 for the incoming rCTP is shown as a green mesh. **(b)** Two focused views of the hydrogen bonds formed between Rev1 R324 and the incoming rCTP. Hydrogen bonds are shown as black dashed lines and hydrogen bonding distances (Å) are labeled. **(c)** A focused view of the incoming nucleotide position in comparison to the Rev1 F367 steric gate residue in the Rev1/DNA/rCTP (green sticks, top) and Rev1/DNA/dCTP (grey sticks, bottom) ternary complex structures. Key distances (Å) are shown as black dashed lines. The 2′-OH in the Rev1/DNA/dCTP complex is modeled. **(d)** An overlay of the Rev1/DNA/rCTP ternary complex (green sticks) and Rev1/DNA/dCTP ternary complex (grey sticks). Red arrows and the respective distances (Å) highlight key active site differences between the two structures.

The change in the rCTP sugar pucker prevents the 2’-OH from clashing with the Rev1 steric gate. However, comparison of the Rev1-rCTP and Rev1-dCTP ternary complex structures revealed several additional conformation changes in the Rev1 active site that likely effect catalysis (Fig. 6d). The C2’-endo sugar pucker of the rCTP results in a 1.3 Å downward movement in the ribose sugar towards the primer terminal dG, which results in a 1.6 Å movement of the primer terminal dG away from the rCTP. In this conformation, the 3’-OH of the primer terminal dG is 5.4 Å away from the Pα of the rCTP and not in proper position for the in-line nucleophilic attack. Therefore, the Rev-rCTP ternary complex structure provides a rational for comparable dCTP and rCTP binding to Rev1, but also explains the decrease in catalysis during Rev1-mediated insertion of rCTP by alterations at the primer terminal 3’-OH^37^.

### Dynamics of the Rev1-rCTP ternary complex active site from MD simulations

Our structural analysis indicates that the incoming rCTP can form a stable interaction with the Rev1 R324 protein-template, and that accommodation of rCTP in the active site is mediated by alteration to the ribose sugar pucker. To obtain additional insight into the stability and dynamics of the R324-rCTP interaction and the rCTP sugar pucker, a 1 μs MD simulation was performed. During the Rev1-rCTP ternary complex simulation, the O2 an N3 of the rCTP Watson-Crick edge and the Nε and Nη were observed within hydrogen bonding distance for 100% and 99.9% of snapshots of the MD trajectory, respectively (Fig. 7a). This indicates the rCTP forms a stable interaction with R324, which is similar to what was observed in the Rev1-dCTP ternary complex structure (Supplementary Fig. 4). Next, to understand the stability and dynamics of the rCTP sugar pucker, the pseudorotation angle of the rCTP ribose sugar was calculated throughout the MD simulation, which reports on the sugar pucker of the rCTP ribose. The pseudorotation angle of the rCTP sugar throughout the MD trajectory ranged from 135-227 with the average being 186.3, indicating the rCTP largely adopts a C2’-endo sugar pucker in the Rev1 active site (Fig. 7b). In contrast, the dCTP ribose largely adopted a C3’-endo sugar pucker (Fig. 7c). However, infrequent sampling of the C2’-endo sugar pucker was observed for dCTP during the simulation. This indicates that the C2’-endo sugar pucker is important for stabilizing the rCTP in the Rev1 active site, and likely explains the comparable affinity of Rev1 for rCTP and dCTP^36, 37^.

**Figure 7.**
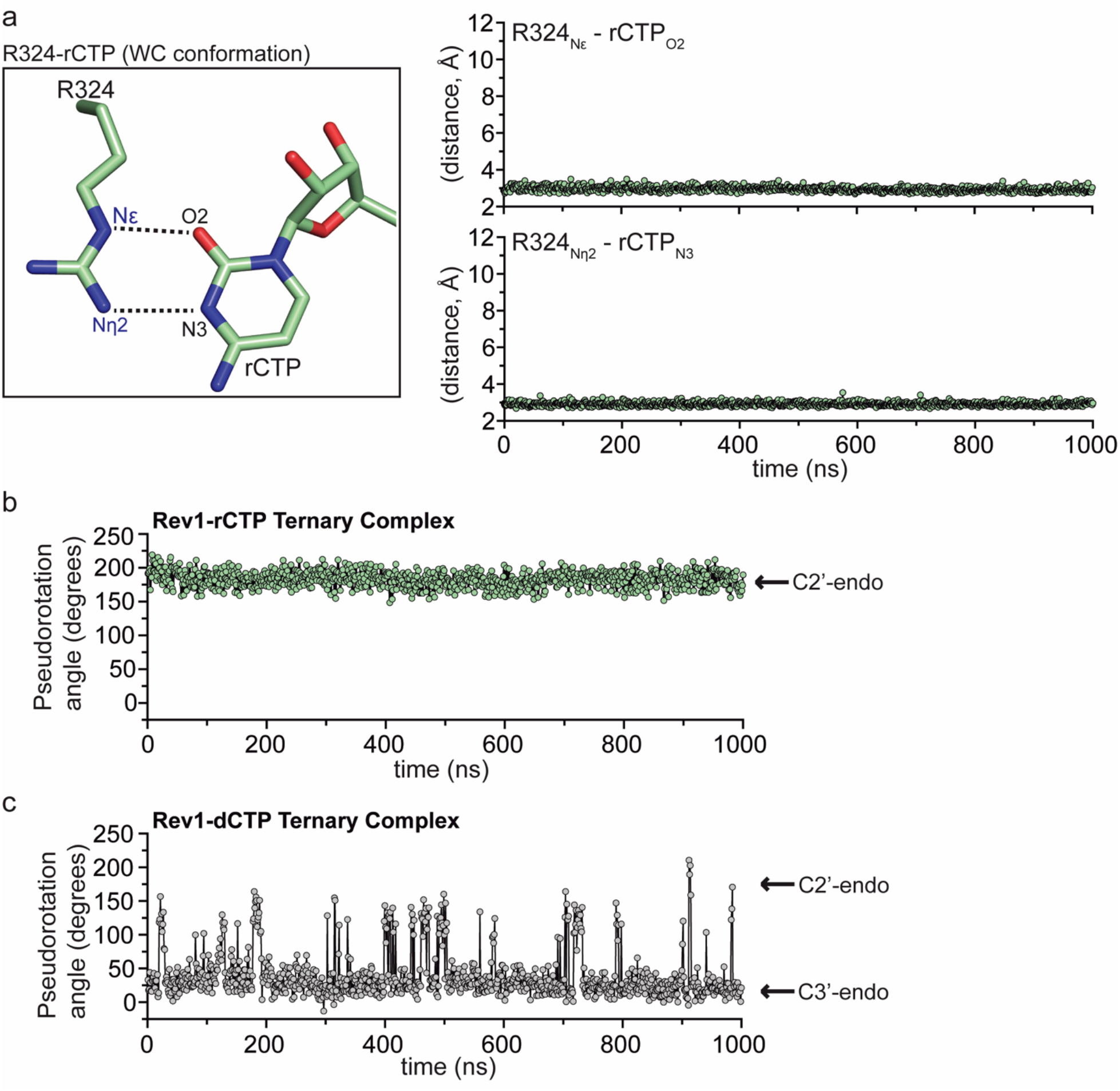
Molecular dynamics simulation of Rev1/DNA/rCTP ternary complex structure. **(a)** Focused view of the Rev1 R324 and incoming rCTP (Watson-crick conformation) showing the distances monitored throughout the MD simulation (left). Distance profiles for R324_NE_-rCTP_O2_ and the R324_Nn2_-rCTP_N3_ in the Rev1/DNA/rCTP ternary complex simulation (right). Each datapoint represents the distance (Å) between the indicated atoms at a single snapshot (1ns) from the MD simulation. **(b)** Pseudorotation angle profiles for the rCTP ribose sugar. Each datapoint represents the pseudorotation angle of the rCTP ribose sugar at a single snapshot (1ns) from the MD simulation. **(c)** Pseudorotation angle profiles for the dCTP ribose sugar. Each datapoint represents the pseudorotation angle of the dCTP ribose sugar at a single snapshot (1ns) from the MD simulation.

## Discussion

### Insight into Rev1 base fidelity

Rev1 utilizes a protein-template mechanism for DNA base discrimination, where the side chain of R324 forms two stabilizing hydrogen bonds with the incoming dCTP. Notably, similar stabilizing hydrogen bonds are unable to be formed with incoming dTTP, dGTP, or dATP, which has led to the hypothesis that R324 may act as a gate to prevent binding of mismatched incoming nucleotides. Notably, this hypothesis has proven difficult to address in the absence of high-resolution structures of Rev1 in complex with mismatched incoming dNTPs. Our work has provided novel mechanistic insight into how Rev1 utilizes the R324 protein template to prevent optimal hydrogen bonding with incoming dTTP, dGTP, and dATP, thus accounting for the enzyme’s substrate specificity. The Rev1-dTTP ternary complex structure indicates that incoming dTTP binding is accomplished through a single non-planar hydrogen bond with R324, which is different than the two hydrogen bonds seen for incoming dCTP (Fig. 2). The non-planar interaction of dTTP and R324 results in a conformation of the 3′-OH of the primer terminal nucleotide that is not in proper orientation for catalysis. These structural changes explain the ∼10-fold decrease in dissociation constant (K_d_) for dTTP binding and ∼26-fold decrease in nucleotide incorporation (k_pol_) previously determined for Rev1 insertion of dTTP using pre-steady state kinetics methods^37^. Next, the Rev1-dGTP ternary complex structure shows that incoming dGTP adopts a Hoogsteen conformation, which allows the dGTP to form two non-planar hydrogen bonds with R324 (Fig. 3). In this conformation, the dGTP is readily accommodated in the spacious Rev1 active site, and results in minimal changes to the 3′-OH of the primer terminal nucleotide. The structural changes to dGTP and minimal changes to the primer terminal 3′-OH are consistent with the ∼45-fold decrease in dissociation constant for dGTP binding and a modest ∼3.5-fold decrease in nucleotide incorporation. Finally, the Rev1-dATP ternary complex structure indicates the dATP is dynamic adopting two distinct conformations in the Rev1 active site. In both conformations, the dATP is in the Hoogsteen conformation and forms a single non-planar hydrogen bond with R324 (Fig. 4).

These non-planar interactions lead to a conformation of the 3′-OH of the primer terminal nucleotide that is not in proper orientation for catalysis. These structural changes explain the ∼30-fold decrease in dissociation constant for dATP binding and a ∼450-fold decrease in nucleotide incorporation for Rev1 insertion of dATP.

Substantial evidence indicates that DNA polymerases select and incorporate the correct dNTP using an induced-fit mechanism, where the DNA polymerase undergoes conformational changes during dNTP binding and sampling of proper Watson-crick base pairing with the templating DNA base^4–8^. We previously showed Rev1 does not undergo a conformational change upon incoming nucleotide binding^34^, nor does Rev1 use a templating DNA base for coordination of the incoming nucleotide during the catalytic cycle^29–31, 33^. The Rev1 ternary complex structures described here highlight how fidelity is maintained through sub-optimal hydrogen bonds of dTTP, dGTP, and dATP with the R324 protein-template. Importantly, this is analogous to the sub-optimal hydrogen bonds between mismatched dNTPs and templating DNA bases for other DNA polymerases. In addition, we identified conformational changes to the primer terminal DNA base that result in a Rev1 active site where the 3’-OH is not in position for the in-line nucleophilic attack on the Pα of the incoming dNTP. This is consistent with the other DNA polymerases, where the 3’-OH is not properly aligned with Pα of the incoming dNTP during insertion of mismatched nucleotides^10, 38–41^. Consistently, our MD simulations suggest that the sub-optimal hydrogen bonds with R324 result in a higher conformational flexibility of the incoming dTTP, dGTP, and dATP in the Rev1 active site. Together, this suggests that optimal hydrogen bonding with the incoming nucleotide and stringent active site organization of the primer terminal 3’-OH are mis-insertion checkpoint used by all DNA polymerases, irrespective of whether a templating DNA base is utilized.

### Insight into Rev1 sugar fidelity

In addition to DNA base discrimination, DNA polymerases must also choose the incoming nucleotide with the correct sugar. This is particularly challenging given that the cellular concentration of ribonucleotides are in vast excess of deoxyribonucleotides^42, 43^. For most DNA polymerases, ribonucleotide discrimination is accomplished through a bulky aromatic steric gate residue, which prevents efficient binding of rNTPs via steric clashes with the 2′-OH^11, 12^. In some cases, the steric gate can also be supplemented with a polar filter, which interacts with the 3′ -OH and brings the C2′ in close proximity to the steric gate^13^. Interestingly, as shown by pre-steady state kinetics, hRev1 binds rCTP with a similar Kd as dCTP, but has a 280-fold decrease in catalysis, indicating that Rev1 sugar discrimination occurs at the catalytic step and not nucleotide binding. The modest sugar fidelity of Rev1 has been perplexing, as Rev1 possesses both a steric gate (F367) and polar filter (S210) residue. Our Rev1-rCTP ternary complex structure and MD simulations provide a rationale for this observation. In the Rev1 active site, the rCTP adopts a C2′-endo sugar pucker that prevents significant clashes with the F367 steric gate (Fig. 6). In addition, the conformation observed for the rCTP is incapable of forming an interaction with the S210 polar filter. This indicates the C2’-endo sugar pucker prevents the incoming rCTP from clashing with the steric gate and polar filter active site residues, and ultimately explains the similar affinity of Rev1 for incoming rCTP and dCTP. Although Rev1 interacts with rCTP and dCTP with similar affinity, a decrease in catalysis for rCTP has been observed^37^. Our structures indicate the decrease in catalysis during Rev1 insertion of rCTP is due to a conformational change in the 3′-OH of the primer terminal DNA base, which arises from the rCTP adopting a 2′-endo sugar pucker in the Rev1 active site. This prevents proper Rev1 active site organization for phosphodiester bond formation between the 3’-OH and the Pα of the incoming nucleotide, which is similar to observations for DNA Pol β and Pol η during mis-insertion of ribonucleotides^44, 45^. Notably, the mechanism for ribonucleotide discrimination by Rev1 is different than that observed for Pol μ and Pol λ, underscoring the importance of studying mechanisms of ribonucleotide discrimination for different DNA polymerases^46, 47^.

### Biological Implications for Rev1 DNA base and sugar fidelity

Our structural studies have provided novel insight into how the R324 protein-template enables Rev1 to maintain preferential insertion of dCTP. Importantly, the preferential insertion of dCTP by Rev1 is advantageous during multiple cellular processes. For example, Rev1 accommodates several forms of adducted guanine bases in its active site, which is important for DNA damage bypass during TLS^30, 31, 33^. In addition, Rev1 is known to perform DNA synthesis across from G-quadruplex sequences during DNA replication^15, 16, 48^. In these biological contexts, it is critical for Rev1 to maintain preferential insertion of dCTP to ensure maintenance of the DNA coding potential. More recently, Rev1 has also been implicated in replication-induced gap filling, which may be a vulnerability in cancer cells ^17, 49^. Consistently, inhibition of Rev1 function has shown promise as a chemotherapeutic adjuvant^50–52^, underscoring the importance for further characterizing the molecular basis of Rev1 function.

The cellular concentrations of rCTP are in vast excess of dCTP^42^. Given the modest sugar fidelity of Rev1 *in vitro*, dCTP and rCTP are likely inserted at similar efficiencies during TLS *in vivo*^37^. Interestingly, the TLS polymerases Pol η and Pol ι also maintain modest sugar selectivity compared to replicative DNA polymerases *in-vitro*^53, 54^. The poor sugar fidelity by TLS polymerases suggests that DNA damage bypass during replication may contribute a small proportion of ribonucleotides found embedded in the genome. Interestingly, RNase H2 is the enzyme responsible for processing ribonucleotides embedded in DNA, and is known to interact with the core TLS component PCNA at stalled replication forks^55–58^. The interaction of PCNA and RNase H2 may allow for the rapid removal of mis-inserted ribonucleotides after DNA damage bypass by the ribonucleotide excision repair (RER) pathway^43^. However, recent evidence suggests that RNase H2 activity is decreased when removing ribonucleotides across from DNA damage^53^, indicating that removal of ribonucleotides after DNA damage bypass by TLS polymerases may be inefficient. It is interesting to speculate whether TLS polymerases may contribute to the accumulation of ribonucleotides in the genome, though additional work is needed to understand the complex interplay between ribonucleotide mis-incorporation during TLS and RNase H2-dependent RER pathways.

## Methods

### Preparation of DNA substrates

For X-ray crystallography, oligonucleotides 5′-ATC-GCT-ACC-ACA-CCC-CT-3′ (template strand) and 5′-GGG-GTG-TGG-TAG-3′ (primer strand) were resuspended in a buffer containing 10 mM Tris-8.0 and 1mM EDTA and annealed by heating to 90 °C for 5 minutes before cooling to 4 °C using a linear gradient (-1 °C min^-1^).

### Expression and Purification of Rev1

The construct for the yeast Rev1 catalytic core was purchased from GenScript. All Rev1 constructs were transformed and expressed in BL21(DE3) plysS E. coli cells (Invitrogen). Cells were grown at 37 °C until an OD_600_ ∼0.7 was reached. Rev1 protein expression was induced with 0.1 mM IPTG at 20 °C overnight. The resulting cell pellets were frozen and stored at -20 °C, or - 80 °C for longer term storage. For cell lysis, Rev1 cell pellets were resuspended in a lysis buffer containing 50 mM HEPES (pH-7.4), 150 mM NaCl, 1 mM EDTA, 1 mM DTT and a cocktail of protease inhibitors. The cells were lysed via sonication and the lysate cleared at 24,000 xg for one hour. The supernatant containing GST-Rev1 protein was incubated with glutathione agarose resin (Goldbio) for 2 hours, washed on the glutathione resin with a high salt buffer containing 50 mM HEPES and 1M NaCl, and the GST-tag cleaved off overnight using PreScission Protease. The cleaved Rev1 protein was purified by cation-exchange chromatography using a POROS HS column (GE Health Sciences) and gel filtration using a HiPrep 16/60 Sephacryl S-200 HR (GE Health Sciences). The purified Rev1 protein was frozen and stored at -80 °C in a buffer containing 250 mM NaCl, 50 mM Tris (pH-8.0), and 2mM TCEP.

### X-ray Crystallography

Oligonucleotides 5′-ATC-GCT-ACC-ACA-CCC-CT-3′ (template strand) and 5′-GGG-GTG-TGG-TAG-3′ (primer strand) were utilized for all X-ray crystallography experiments. The annealed DNA substrate (1.15 mM) was mixed with Rev1 protein (5-6 mg ml^-1^) and the Rev1/DNA binary crystals obtained in a condition with the reservoir containing 15-23% PEG3350 and 200 mM ammonium nitrate using the sitting-drop vapor diffusion method. To generate the dTTP and rCTP ternary complex structures, the Rev1/DNA binary crystals were transferred to a cryoprotectant containing reservoir solution, 25% glycerol, 50 mM CaCl_2_ and 10 mM of dTTP or rCTP. Attempts to soak in dATP and dGTP into the Rev1 active site were unsuccessful. To generate dATP and dGTP ternary complex structures, the Rev1 protein (5-6 mg ml^-1^), annealed DNA substrate (1.15 mM), 50 mM CaCl_2_, and dATP (1 mM) or dGTP (20 mM) were mixed and crystals obtained in a condition with the reservoir containing 15-23% PEG3350 and 200 mM ammonium nitrate using the sitting-drop vapor diffusion method. The dATP and dGTP ternary complex crystals were transferred to a cryoprotectant containing reservoir solution and 25% glycerol. All X-ray crystallography data were collected at 100 K on a Rigaku MicroMax-007 HF rotating anode diffractometer equipped with a Dectris Pilatus3R 200K-A detector system at a wavelength of 1.54 Å. Initial models were generated by molecular replacement using a previously solved Rev1/DNA/dCTP ternary complex (PDB: 6X6Z or 5WM1) as a reference structure. Further refinement and model building was carried out using PHENIX and Coot, respectively^59, 60^. All figures containing structures were generated using PyMOL (Schrödinger LLC)^61^. The sugar puckers for incoming NTPs were determined by calculating pseudorotation angles using the web 3DNA server^62^.

### MD Simulations

Molecular dynamics (MD) simulations were performed for Rev1 and DNA complexes in explicit water solvent, using a protocol similar to our previous study^34^. Model preparation and simulations were performed using the AMBER v16 suite of programs for biomolecular simulations^63^. AMBER’s *ff14SB* ^64^ force-fields were used for all simulations. MD simulations were performed using NVIDIA graphical processing units (GPUs) and AMBER’s *pmemd.cuda* simulation engine using our lab protocols published previously^65, 66^.

A total of 5 separate simulations were performed for Rev1-dTTP, Rev1-dGTP, Rev1-dATP (in Hoogsteen conformation, conformation 2) Rev1-dATP (in Watson-Crick conformation) and Rev1-rCTP (all as ternary complexes) based on the X-ray crystal structures determined in this study. The missing hydrogen atoms were added by AMBER’s *tleap* program. After processing the coordinates of the protein and substrate, all systems were neutralized by addition of counter-ions and the resulting system were solvated in a rectangular box of SPC/E water, with a 10 Å minimum distance between the protein and the edge of the periodic box. The prepared systems were equilibrated using a protocol described previously ^67, 68^. The equilibrated systems were then used to run 1.0 μs of production MD under constant energy conditions (NVE ensemble). The use of NVE ensemble is preferred as it offers better computational stability and performance ^69, 70^. The production simulations were performed at a temperature of 300 K. As NVE ensemble was used for production runs, these values correspond to initial temperature at start of simulations. Temperature adjusting thermostat was not used in simulations; over the course of 1.0 μs simulations the temperature fluctuated around 300 K with RMS fluctuations between 2-4 K, which is typical for well equilibrated systems. A total of 1,000 conformational snapshots (stored every 1,000 ps) collected for each system was used for analysis.

Distance profile calculations: The distances and percentage occupancy were calculated using 1,000 stored during the MD simulations. A cutoff of 3.5 Å was used to calculate the percent of snapshots within hydrogen bonding distance (occupancy) during each MD simulation.

## Acknowledgments

We thank Nicole Hoitsma (University of Kansas Medical Center) for helpful discussion and assistance with structural modeling. This research was supported by the National Institute of General Medical Science [R35-GM128562 to B.D.F., T.M.W, T.H.K.] and [R01 - GM081433 to M.T.W].

## Data Availability

Atomic coordinates and structure factors for the reported crystal structures have been deposited with the Protein Data bank under accession numbers 7T18, 7T19, 7T1A, and 7T1B.

---

**Supplementary Figure 1.**
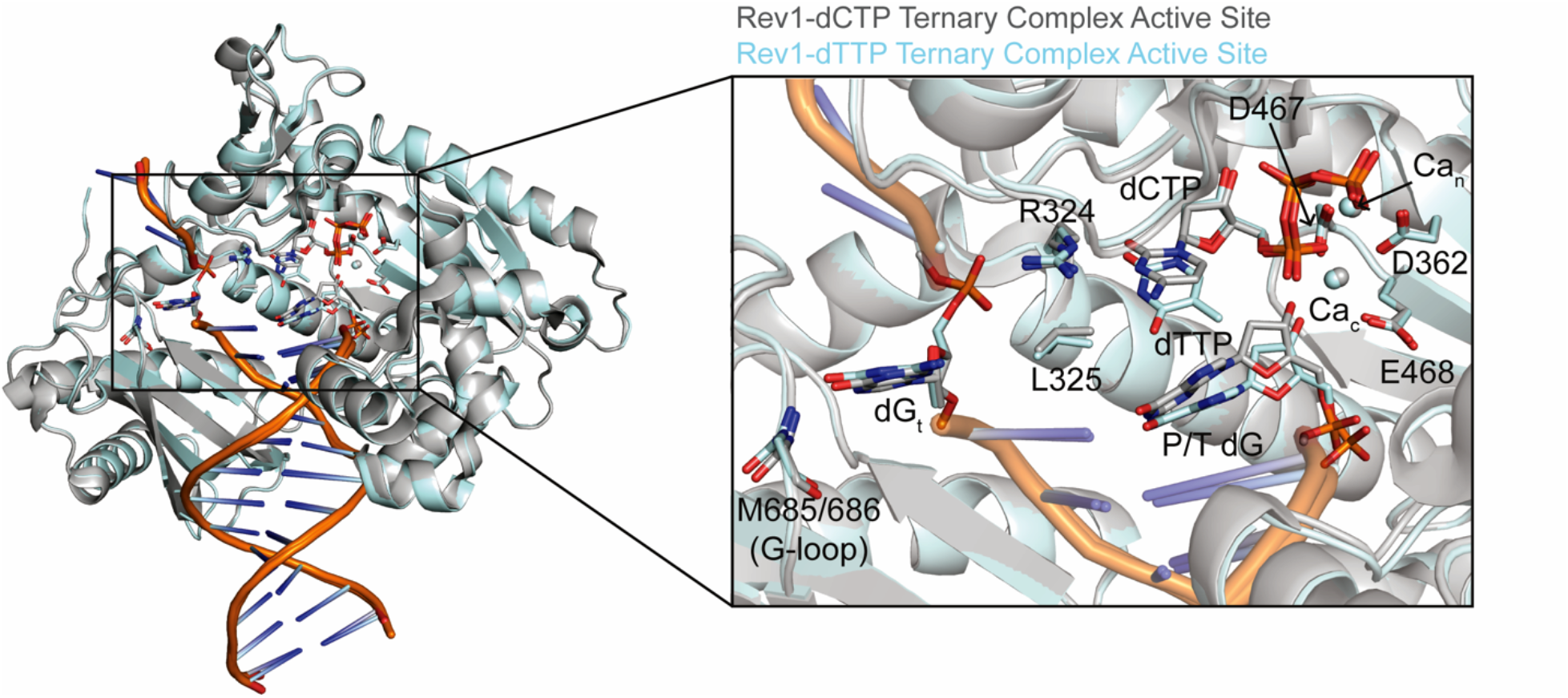
Overall comparison of the Rev1/DNA/dTTP and Rev1/DNA/dCTP ternary complex structures. Comparison of the overall structure (left) and active site (right) of the Rev1/DNA/dTTP (blue) and Rev1/DNA/dCTP (grey) ternary complexes. Key protein and DNA residues are shown as sticks.

**Supplementary Figure 2.**
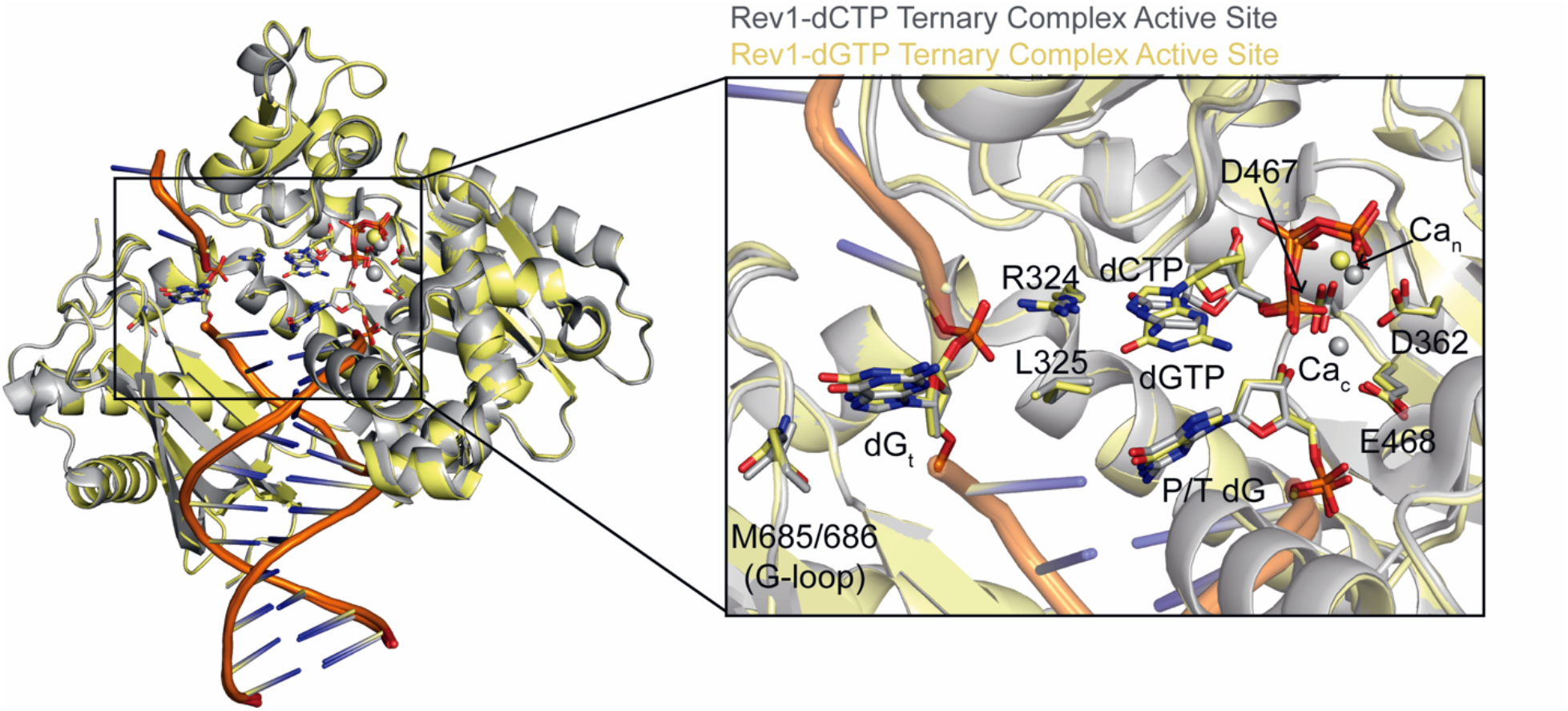
Overall comparison of the Rev1/DNA/dGTP and Rev1/DNA/dCTP ternary complex structures. Comparison of the overall structure (left) and active site (right) of the Rev1/DNA/dGTP (yellow) and Rev1/DNA/dCTP (grey) ternary complexes. Key protein and DNA residues are shown as sticks.

**Supplementary Figure 3.**
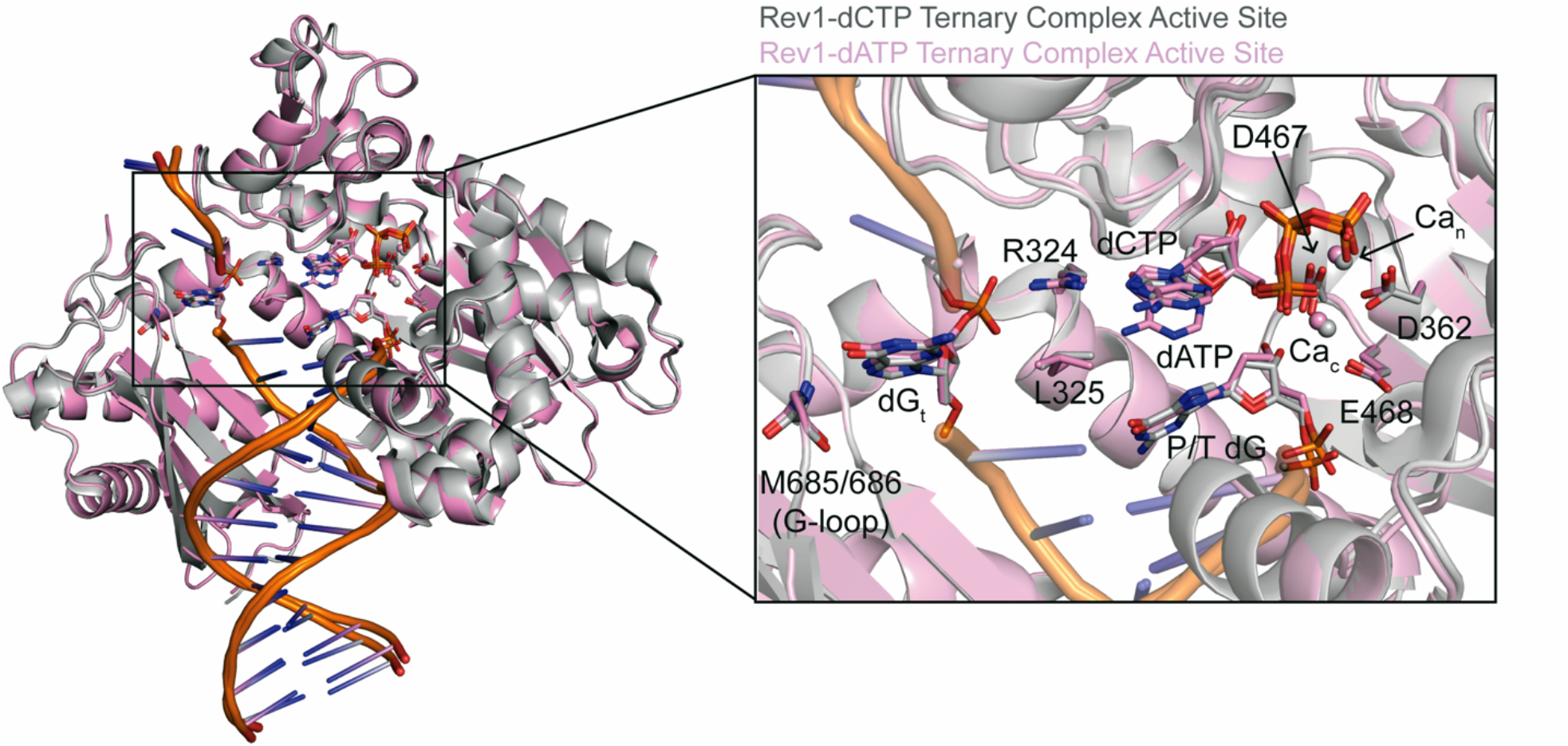
Overall comparison of the Rev1/DNA/dATP and Rev1/DNA/dCTP ternary complex structures. Comparison of the overall structure (left) and active site (right) of the Rev1/DNA/dATP (pink) and Rev1/DNA/dCTP (grey) ternary complexes. Key protein and DNA residues are shown as sticks.

**Supplementary Figure 4.**
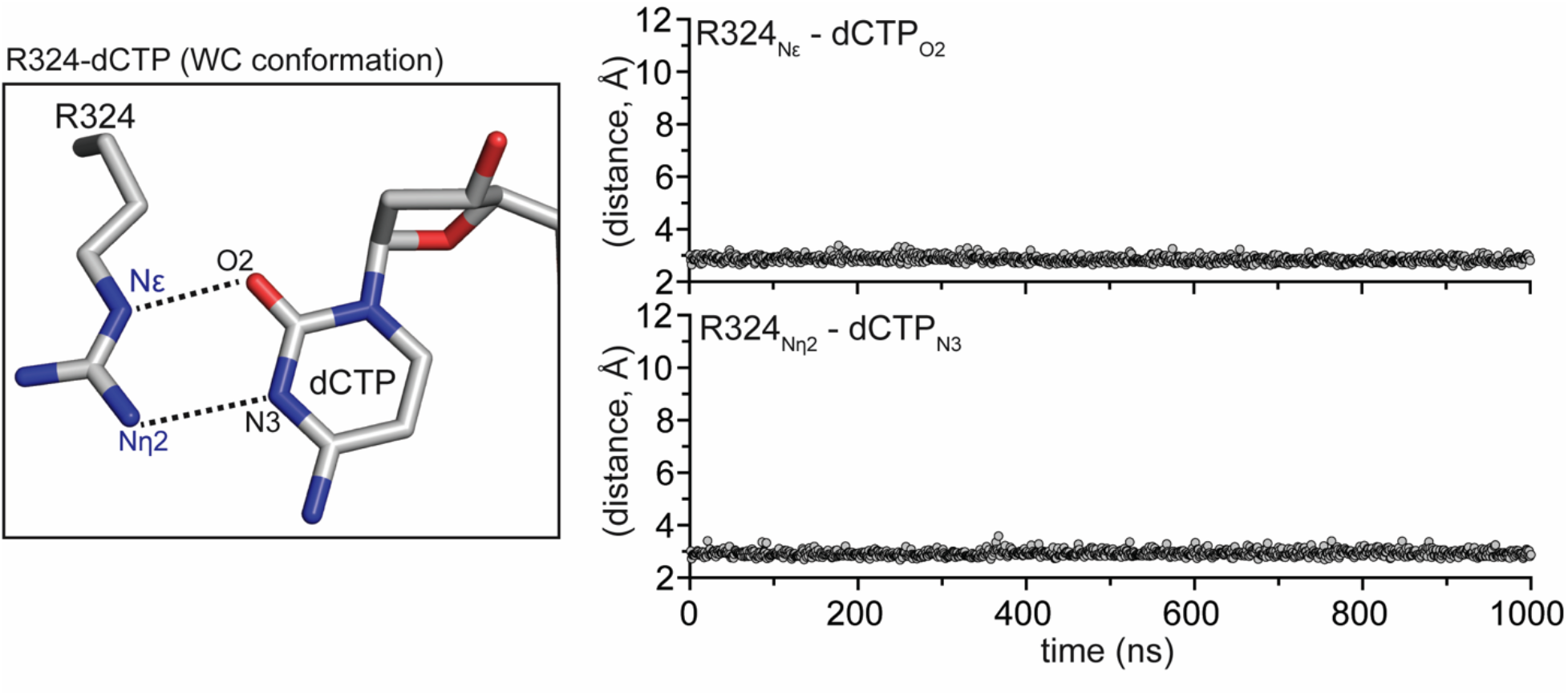
Molecular dynamics simulation of Rev1/DNA/dCTP ternary complex structure. Focused view of the Rev1 R324 and incoming dCTP (Watson-crick conformation) showing the distances monitored throughout the MD simulation (left). Distance profiles for R324_NE_-dCTP_O2_ and the R324_Nn2_-dCTP_N3_ in the Rev1/DNA/dCTP ternary complex simulation (right). Each datapoint represents the distance (Å) between the indicated atoms at a single snapshot (1ns) from the MD simulation.

**Supplementary Figure 5.**
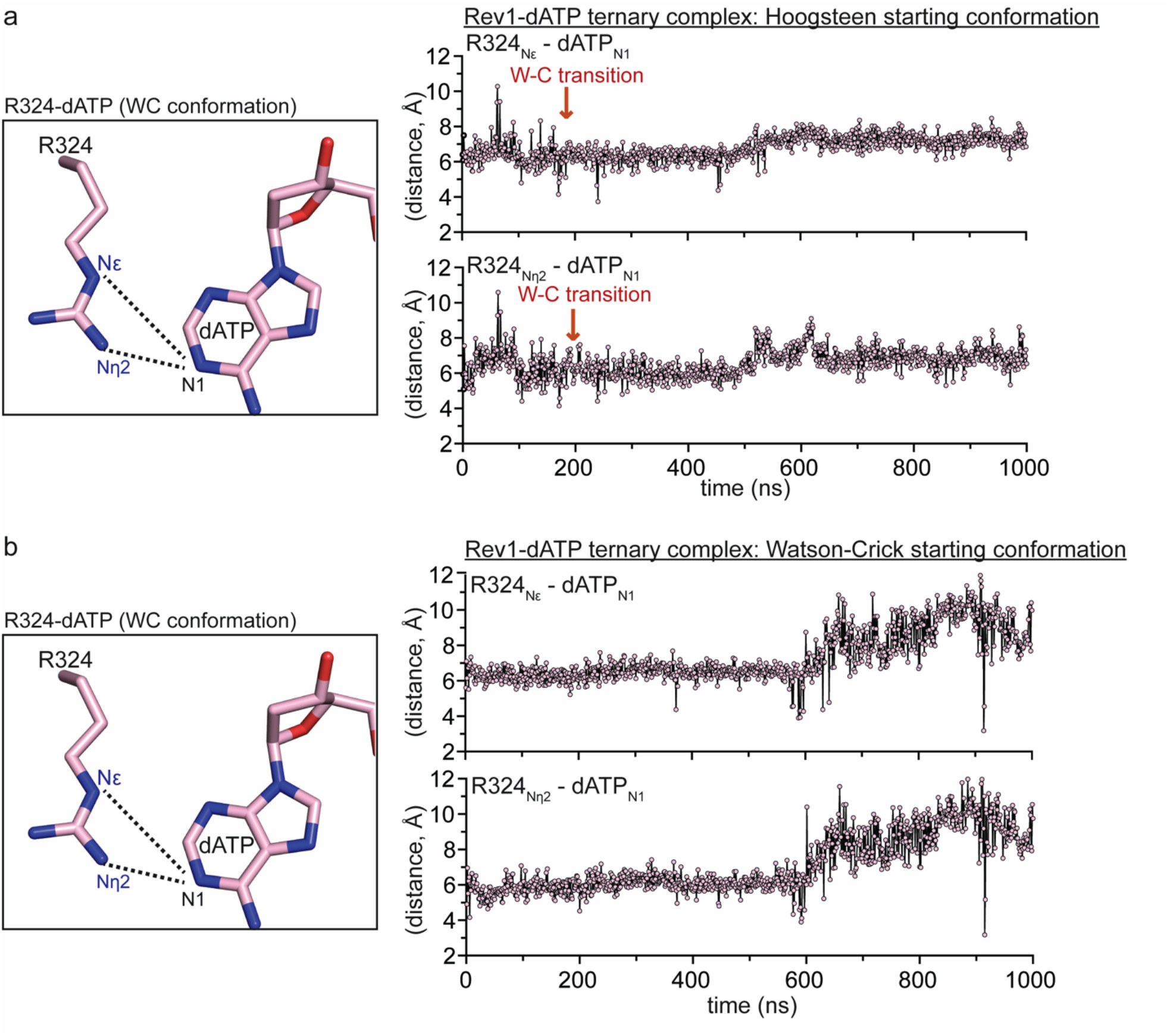
Molecular dynamics simulation of Rev1/DNA/dATP ternary complex structure. **(a)** Focused view of the Rev1 R324 and incoming dATP (Watson-crick conformation) showing the distances monitored throughout the MD simulation (left). Distance profiles for R324_NE_-dATP_N1_ and the R324_Nn2_-dATP_N1_ in the Rev1/DNA/dATP ternary complex simulation. Each datapoint represents the distance (Å) between the indicated atoms at a single snapshot (1ns) from the MD simulation. The MD simulation was started with the dATP in the Hoogsteen conformation. The dATP transition from Hoogsteen to Watson-Crick conformation is denoted with a red arrow. **(b)** Focused view of the Rev1 R324 and incoming dATP (Watson-crick conformation) showing the distances monitored throughout the MD simulation (left). Distance profiles for R324_NE_-dATP_N1_ and the R324_Nn2_-dATP_N1_ in the Rev1/DNA/dATP ternary complex simulation. Each datapoint represents the distance (Å) between the indicated atoms at a single snapshot (1ns) from the MD simulation. The MD simulation was started with the dATP in the theoretical Watson-crick conformation.

**Supplementary Figure 6.**
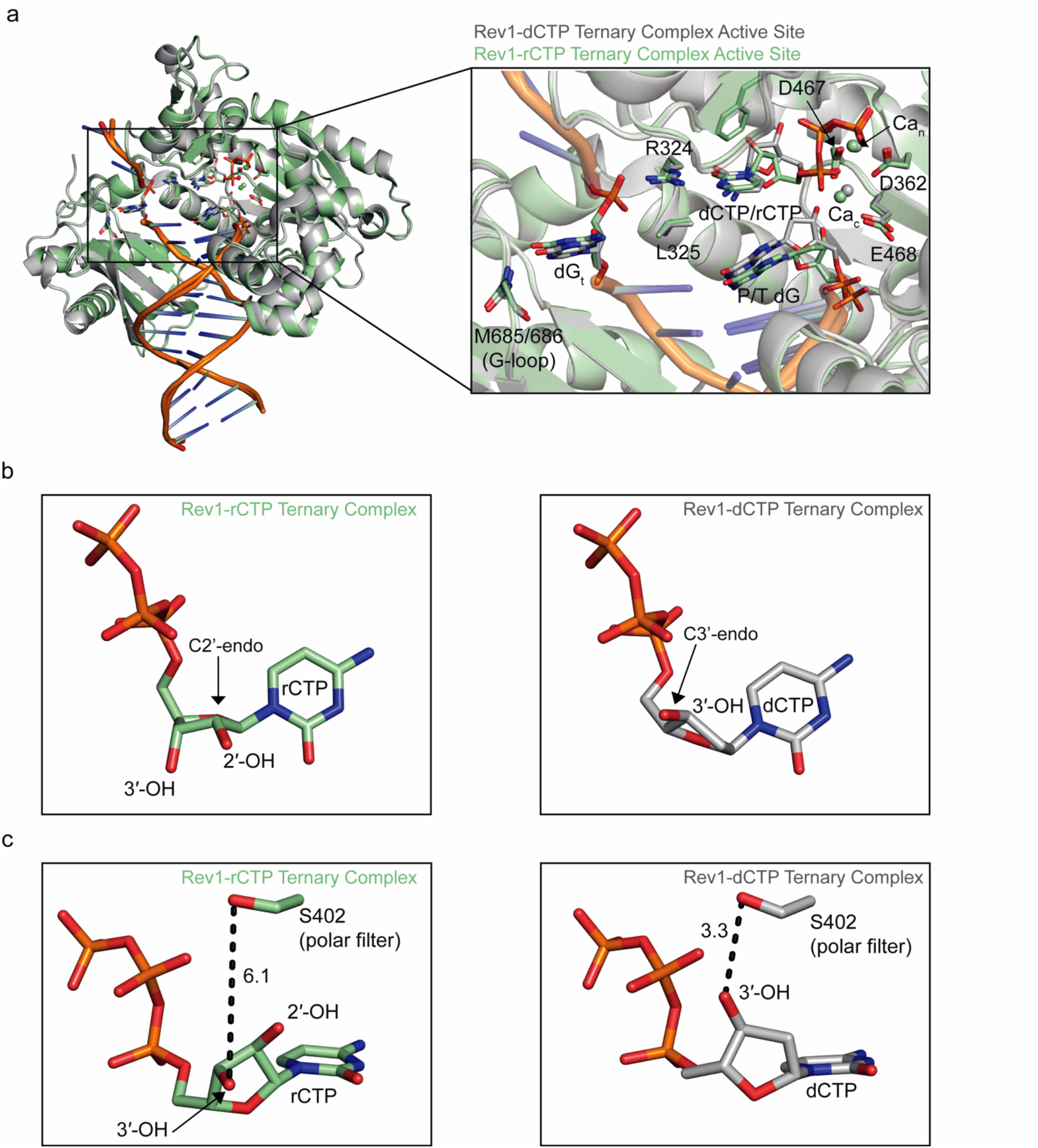
Overall comparison of the Rev1/DNA/rCTP and Rev1/DNA/dCTP ternary complex structures. **(a)** Comparison of the overall structure (left) and active site (right) of the Rev1/DNA/rCTP (green) and Rev1/DNA/dCTP (grey) ternary complexes. Key protein and DNA residues are shown as sticks. **(b)** A focused view of the incoming nucleotide sugar pucker in the Rev1/DNA/rCTP (left) and Rev1/DNA/dCTP (right) ternary complex structures. **(c)** A focused view of the incoming nucleotide position and Rev1 S402 polar filter residue in the Rev1/DNA/rCTP (green sticks, left) and Rev1/DNA/dCTP (grey sticks, right) ternary complex structures. Key distances (Å) are shown as black dashed lines.

**Supplementary Table 1:** Rev1 pre-steady state kinetics c site is a cognate lesion for the yeast Rev1 protein. *DNA repair*, **10**, 1138-1144.

| Organism | NTP | $k_{\text{pol}}$ | $K_d$ (NTP) | $F_{\text{rel}}^1$ |
| --- | --- | --- | --- | --- |
| *H. sapien*<sup>3</sup> | dCTP | $22.4 \pm 0.9$ | $2.2 \pm 0.3$ | - |
| | dTTP | $0.88 \pm 0.06$ | $22.7 \pm 7$ | 260 |
| | dGTP | $6.3 \pm 0.3$ | $90 \pm 10$ | 140 |
| | dATP | $0.050 \pm 0.004$ | $70 \pm 20$ | 14000 |
| | rCTP | $0.098 \pm 0.002$ | $2.7 \pm 0.2$ | 280 |
| *S. cerevisiae*<sup>4</sup> | dCTP | $18 \pm 1$ | $32 \pm 7$ | -- |
| | dTTP | $0.20 \pm 0.02$ | $700 \pm 100$ | 2000 |
| | dGTP | $0.97 \pm .0.04$ | $460 \pm 40$ | 270 |
|  | dATP | ND<sup>2</sup> | ND<sup>2</sup> | ND<sup>2</sup> |

<sup>1</sup> $F_{\text{rel}}$ is the relative catalytic efficiency of incorporating the given dNTP versus dCTP, which is $k_{\text{pol}}(\text{dCTP})/k_{\text{pol}}(\text{dNTP})$ times $K_d(\text{dNTP})/K_d(\text{dCTP})$ .
<sup>2</sup>ND is not detected.
<sup>3</sup>Values from Brown, J.A., Fowler, J.D. and Suo, Z. (2010) Kinetic basis of nucleotide selection employed by a protein template-dependent DNA polymerase. *Biochemistry*, **49**, 5504-5510.
<sup>4</sup>Values from Pryor, J.M. and Washington, M.T. (2011) Pre-steady state kinetic studies show that an abasic site is a cognate lesion for the yeast Rev1 protein. *DNA repair*, **10**, 1138-1144.

**Supplementary Table 2:** Data collection and refinement statistics for Rev1 crystal structures

|  | Rev1/DNA/dTTP<br>Ternary Complex | Rev1/DNA/dGTP<br>Ternary Complex | Rev1/DNA/dATP<br>Ternary Complex | Rev1/DNA/rCTP<br>Ternary Complex |
| --- | --- | --- | --- | --- |
| **Data collection** |  |  |  |  |
| Space group | P2<sub>1</sub>2<sub>1</sub>2<sub>1</sub> | P2<sub>1</sub>2<sub>1</sub>2<sub>1</sub> | P2<sub>1</sub>2<sub>1</sub>2<sub>1</sub> | P2<sub>1</sub>2<sub>1</sub>2<sub>1</sub> |
| Cell dimensions<br>*a*, *b*, *c* (Å) | 62.2,73.6,117.8 | 62.2,73.6,117.0 | 62.1,73.3,115.4 | 62.2,73.5,117.7 |
| $\alpha$, $\beta$, $\gamma$ (°) | 90,90,90 | 90,90,90 | 90,90,90 | 90,90,90 |
| Resolution (Å) | 25 – 1.70 | 25 – 2.03 | 25 – 1.80 | 25 – 1.75 |
| $R_{\text{meas}}$<sup>a</sup> (%) | 0.078 (>1) | 0.165 (>1) | 0.077 (0.924) | 0.084 (.952) |
| $I/\sigma I$ | 15.1 (1.3) | 6.9 (5.0) | 12.5 (0.7) | 11.5 (0.81) |
| $cc1/2$<sup>b</sup> | 0.529 | 0.443 | 0.556 | 0.542 |
| Completeness<sup>a</sup> (%) | 99.8 (99.6) | 99.5 (99.2) | 99.3 (98.6) | 98.1 (94.4) |
| Redundancy<sup>a</sup> | 4.3 (3.0) | 4.4 (3.0) | 3.5 (2.4) | 2.5 (1.7) |
| **Refinement** |  |  |  |  |
| Resolution (Å) | 24.66 – 1.70 | 24.59 – 2.01 | 24.65 – 1.81 | 24.63 – 1.75 |
| No. reflections | 58143 | 24002 | 55331 | 77480 |
| $R_{\text{work}}/R_{\text{free}}$ | 18.23/22.53 | 18.12/24.22 | 19.31/22.96 | 18.29/22.81 |
| No. atoms |  |  |  |  |
| Protein | 3527 | 3570 | 3574 | 3551 |
| DNA | 571 | 568 | 569 | 571 |
| Water | 626 | 212 | 306 | 358 |
| B-factors (Å<sup>2</sup>) |  |  |  |  |
| Protein | 18.52 | 32.13 | 24.96 | 22.43 |
| DNA | 25.65 | 40.97 | 38.45 | 31.60 |
| Water | 30.92 | 35.35 | 31.00 | 31.59 |
| R.m.s deviations |  |  |  |  |
| Bond length (Å) | 0.008 | 0.009 | 0.013 | 0.009 |
| Bond angles (°) | 0.991 | 1.053 | 1.278 | 1.055 |
| PDB ID | 7T18 | 7T19 | 7T1A | 7T1B |

<sup>a</sup> Highest resolution shell is shown in parenthesis<sup>b</sup> Highest resolution shell only.

